# Distinguishing pedigree relationships using multi-way identical by descent sharing and sex-specific genetic maps

**DOI:** 10.1101/753343

**Authors:** Ying Qiao, Jens Sannerud, Sayantani Basu-Roy, Caroline Hayward, Amy L. Williams

**Author notes:** These authors contributed equally to this work (Ying Qiao, Jens Sannerud).

## Abstract

The proportion of samples with one or more close relatives in a genetic dataset increases rapidly with sample size, necessitating relatedness modeling and enabling pedigree-based analyses. Despite this, relatives are generally unreported and current inference methods typically detect only the degree of relatedness of sample pairs and not pedigree relationships. We developed CREST, an accurate and fast method that identifies the pedigree relationships of close relatives. CREST utilizes identical by descent (IBD) segments shared between a pair of samples and their mutual relatives, leveraging the fact that sharing rates among these individuals differ across pedigree configurations. Furthermore, CREST exploits the profound differences in sex-specific genetic maps to classify pairs as maternally or paternally related—e.g., paternal half-siblings—using the locations of autosomal IBD segments shared between the pair. In simulated data, CREST correctly classifies 91.5-99.5% of grandparent-grandchild (GP) pairs, 70.5-97.0% of avuncular (AV) pairs, and 79.0-98.0% of half-siblings (HS) pairs compared to PADRE’s rates of 38.5-76.0% of GP, 60.5-92.0% of AV, 73.0-95.0% of HS pairs. Turning to the real 20,032 sample Generation Scotland (GS) dataset, CREST correctly determines the relationship of 99.0% of GP, 85.7% of AV, and 95.0% of HS pairs that have sufficient mutual relative data, completing this analysis in 10.1 CPU hours including IBD detection. CREST’s maternal and paternal relationship inference is also accurate, as it flagged five pairs as incorrectly labeled in the GS pedigrees— three of which we confirmed as mistakes, and two with an uncertain relationship—yielding 99.7% of HS and 93.5% of GP pairs correctly classified.

## Introduction

Modern scale genetic datasets contain tens to hundreds of thousands of samples, sample sizes within which numerous close relatives exist ^1,2^. Characterizing relatives within these samples is essential to avoid spurious signals and to improve power in genetic association studies ^3–5^, but standard models consider only kinship estimates while ignoring the potential for different relationship types to vary in their phenotypic correlation ^6,7^. Moreover, while population genetic studies typically filter close relatives to avoid modeling violations ^8^, such an approach will dramatically reduce sample sizes in large datasets ^1,2^. One way to enable analyses of full study samples is to directly model the transmission of shared haplotypes—i.e., identical by descent (IBD) segments ^9^—using the pedigree structure of each set of relatives, but this requires accurate determination of those pedigrees. And although several approaches exist for inferring pedigrees from genetic data ^10–12^, ambiguities in the true pedigree relationships limit the utility of these methods.

Identifying pedigree relationships is simple for first degree relatives—parent-child (PC) and full sibling pairs— yet distinguishing relatives only one degree more distant, including grandparent-grandchild (GP), avuncular (AV), and half-sibling (HS) pairs remains a challenge. Most methods infer only the degree of relatedness of a pair using either the number and length of pairwise IBD segments ^13^, or the proportion of their genome a pair shares IBD ^14,15^. However, an existing method that leverages these pairwise signals provides limited ability to discriminate among second degree relationships ^16^. Turning to multi-way IBD approaches, a recent method detects aunts/uncles of siblings ^17^, but it requires at least two siblings to work and can only identify their aunts/uncles.

We developed CREST (Classification of RElationShip Types), an approach for inferring sex-specific pedigree relationships that leverages multi-way IBD sharing and sex-specific genetic maps. CREST utilizes multi-way IBD sharing to differentiate relationship types, relying on the fact that a pair of close relatives are expected to share IBD regions with their mutual relatives at different rates depending on that pair’s relationship. For example, consider a mutual relative that is the parent of the genetically older member of a second degree relative pair. Because each meiosis leads to the transmission of half a parent’s DNA, a grandchild will, in expectation, inherit 1/4 of the regions shared IBD between the grandparent and the mutual relative—i.e., the parent of that grandparent. In the case of AV pairs, since two full siblings share equal amounts of IBD with their parent, the child of one sibling—the niece/nephew of the other—is expected to share 1/2 as many sites IBD with her/his grandparent as the aunt/uncle does. Lastly, two half-siblings have equal IBD sharing with their common parent. These same sharing rates arise for many other types of mutual relatives and enable the classification of relationship types. Thus, we derived IBD quantities based on this idea and trained kernel density estimation models (KDEs) to classify these three types of second degree relatives in CREST.

This approach of leveraging IBD sharing with mutual relatives not only determines the pedigree relationship types of second degree relatives, it also identifies the directionality of the relationship, that is, which sample is genetically older (e.g., which is the grandparent or aunt/uncle). In particular, the sample with higher levels of IBD sharing with mutual relatives is most likely to be from an earlier generation. (Other pedigree inference methods similarly identify this information using kinship coefficients ^10,11^.) CREST applies this logic to GP and AV pairs and to PC relatives to detect which sample is the parent. When available, age information unambiguously implies the genetically older sample for direct descendants (PC and GP relationships), but fails for AV pairs since a niece/nephew can be (temporally) older than an aunt/uncle.

Beyond solely classifying close relationship types among individuals, we introduce a model to infer the sex of ungenotyped parents that connect the close relatives to each other, further refining CREST’s inferred pedigree relationships. More specifically, CREST infers the sex of the shared parent of HS pairs and whether a grandparent is related through the grandchild’s father or mother. While the mean amount of DNA shared between HS and GP pairs is unaffected by the sex of this parent, we leverage the substantial differences in male and female genetic maps ^18^ to distinguish between the two possibilities. The signature of male and female recombinations on IBD segments is strikingly different, to such an extent that we use autosomal IBD segments alone to perform inference. This application of sex-specific maps liberates CREST from requiring sex chromosome or mitochondrial data for inference, which may be less precise than recombination-based inference and would impose additional restrictions on the sample pairs to which it can be applied (i.e., in terms of their sexes).

We used a combination of simulated and real pedigree data to evaluate CREST, the latter from the Generation Scotland ^19,20^ (GS) cohort. The GS data consist of 20,032 samples recruited as part of families, and include 848 GP, 6,599 AV, and 381 HS pairs. We also compared CREST’s results to those of PADRE ^21^, a composite likelihood method that infers pedigree structures for two sets of close relatives when members of the sets are also related to each other. PADRE makes use of the relationship between the two sets to distinguish pedigrees and, in the case of our analysis, infer the pedigree relationship of the second degree pairs.

In addition to classifying second degree relatives, the CREST approach has the potential to be applied to more distant relationship types. To demonstrate this, we show that, when using simulated IBD segments that are free of both false positives and negatives and limited to those *>*5 centiMorgans (cM) long, CREST can also distinguish pedigree relationships of third degree relatives. While this is less practical than the results in second degree relatives—wherein we used IBIS ^22^, a new, fast approach for IBD detection that operates on unphased genotype data—future detection methods may yield IBD segments of sufficient quality to make this application feasible, thus further expanding the range of pedigree relationships that CREST detects.

## Methods

CREST takes inferred IBD segments from a set of samples as input and applies a multi-way IBD sharing analysis to classify pedigree relationships among pairs; it also uses the locations of IBD segments in HS and GP pairs to infer whether they are maternally or paternally related. The multi-way IBD analysis calculates ratios from the lengths of IBD regions shared among a given target pair of close relatives and their mutual relatives, as described below. The algorithm then uses KDEs trained on ratios calculated from simulated data to infer the relationship type.

While CREST can use good quality IBD segments inferred by any method, IBIS works well in large samples, with the trade off that it reliably identifies ∼7 cM or longer segments. Our experimental results indicate that use of these long segments suffices for discriminating between second degree relationship types. Still, as either data resources become richer (e.g., through sequencing) or as later methods enable the detection of shorter segments in large SNP array datasets, the quality of CREST’s inference will increase.

Throughout, we refer to IBD regions that two or more samples share on only one haplotype copy as IBD1, and those the individuals share IBD on both chromosomes as IBD2.

### Multi-way IBD sharing ratios

CREST utilizes the amount of IBD sharing between a pair of close relatives *x*_1_ and *x*_2_ and one or more of their mutual relatives to distinguish their relationship. We begin by describing the approach using one mutual relative *y*.

The expected IBD rates we adopt are based on the assumption that each *y* is related to both *x*_1_ and *x*_2_ only through the most recent common ancestor(s) (MRCA) of *x*_1_ and *x*_2_. In this case, all IBD segments shared between *y* and one or both of *x*_1_ and *x*_2_ must have been transmitted by this/these MRCA. (The assumption also excludes inbred cases in which *x*_1_ or *x*_2_ is also related to *y* through another ancestor that the other is not.) For example, if *x*_1_ is the grandparent of *x*_2_, their MRCA is *x*_1_ itself, and if *y* is the half-sibling of *x*_1_, *y* is related to both *x*_1_ and *x*_2_ only through *x*_1_ (via the common parent of *x*_1_ and *y*), so the assumption holds. However, if *y* is the half-sibling of the grandchild *x*_2_, *y* is related to *x*_2_ through their common parent, and not only through *x*_1_, in conflict with the assumption. In fact, mutual relatives that are descendants of either *x*_1_ or *x*_2_ violate the assumption in many cases. To exclude direct descendants of *x*_1_ and *x*_2_, we only analyze mutual relatives that are third degree (e.g., a first cousin) or more distantly related to both *x*_1_ and *x*_2_. Because most genetic datasets only span two or three generations, this strategy should generally prevent analyses involving descendant mutual relatives.

The intuition behind the approach is that *x*_1_ and *x*_2_ will share different relative amounts of IBD regions with *y* depending on their relationship. We use two ratios to quantify the IBD sharing rates:

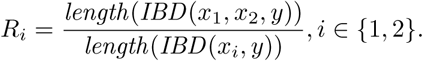

Here *IBD* (*s*_1_, *s*_2_, …, *s*_*n*_) denotes the set of IBD regions that all samples *s*_1_, *s*_2_, …, *s*_*n*_ share, i.e., the intersection of the IBD segments each of the 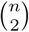 pairs share. The *length* function sums the genetic length (i.e., Morgan [M] length) of a set of IBD segments, accounting for the diploid status of each segment. That is, for a given set of IBD segments *I*,

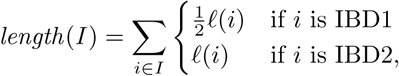

where *ℓ*(*i*) denotes the (M) genetic length of an IBD segment *i* (here from a sex averaged genetic map). The numerators are the same in both ratios and give the genetic length of IBD regions shared jointly by all three samples. The denominators are the length of IBD segments shared by *x*_1_ and *y* in *R*_1_, and by *x*_2_ and *y* in *R*_2_.

These ratios differ according to the relationship type of the second degree relatives. Specifically, for a GP pair, if *x*_1_ is the grandparent of *x*_2_, the numerator *length*(*IBD* (*x*_1_, *x*_2_, *y*)) = *length*(*IBD* (*x*_2_, *y*)) since *x*_2_ will inherit a subset of the IBD segments *x*_1_ shares with *y* (Figure 1A). Additionally, 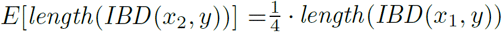 since *x*_2_ is two meioses away from *x*_1_ and each meiosis leads to the transmission of an average of one-half of the IBD segment length any pair of relatives shares. Thus, 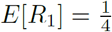 and *E*[*R*_2_] = 1. Similarly, each member of a HS pair independently inherits one-half of the genome of their common parent 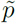, so the probability that they both inherit a given IBD region that 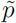 and *y* share is 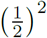 (Figure 1B). Therefore the expected numerator is 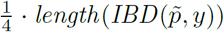, and the expected denominator is 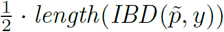 for both *R*_1_ and *R*_2_, so 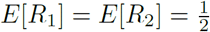. In the case of an AV pair, the aunt/uncle inherits half the genome of her/his parent 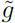—the grandparent of the niece/nephew—that is related to *y*. And as in the GP case, the niece/nephew is expected to inherit one-quarter of the genome of 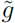 (Figure 1C). Therefore the expected numerator is 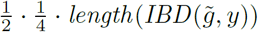, the expected denominator of *R*_1_ is 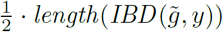, and that of *R*_2_ is 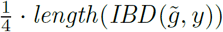, resulting in 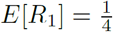 and 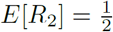.

**Figure 1:**
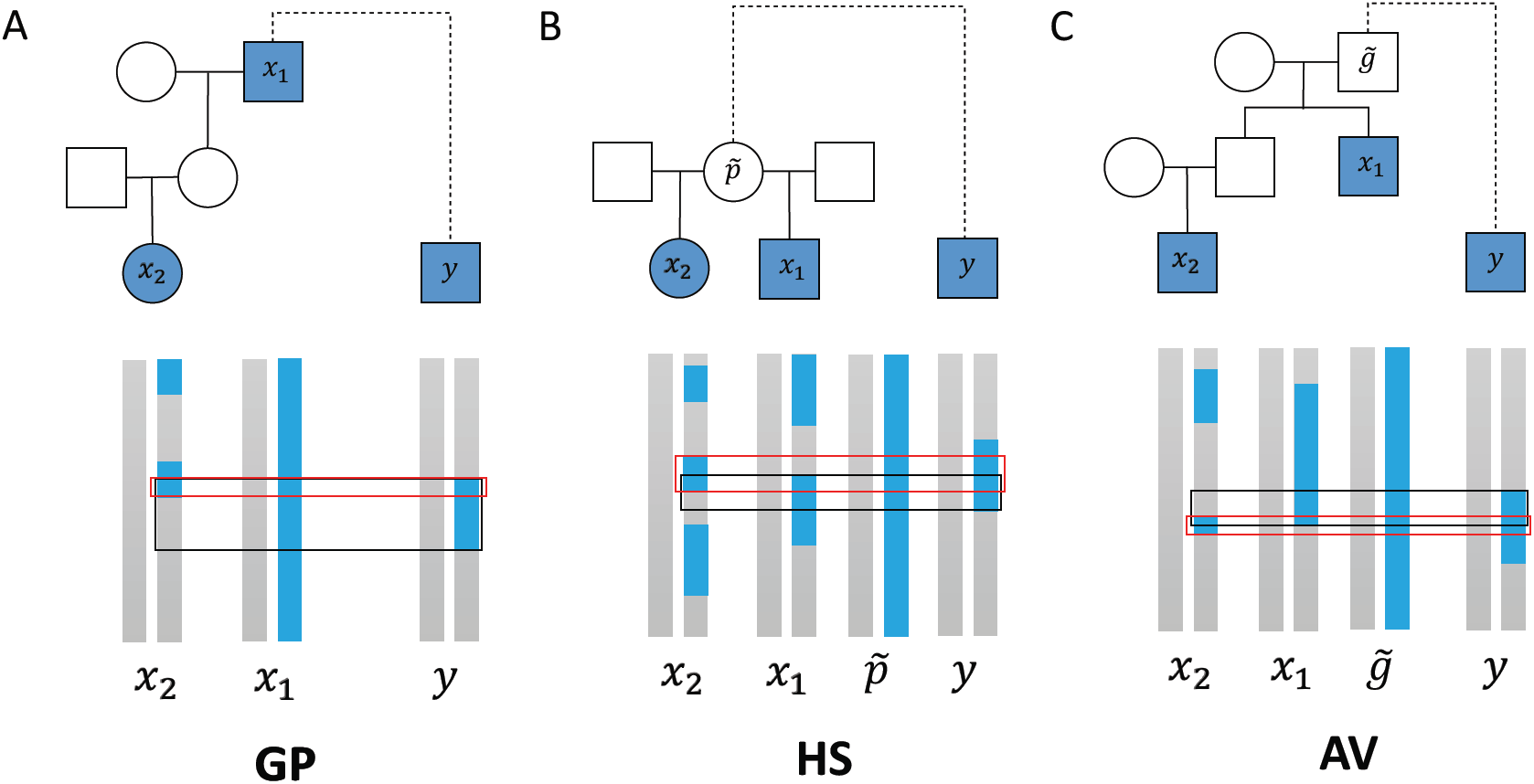
Example IBD sharing between the three types of second degree relatives and one of their mutual relatives. Samples with filled shapes are those for which data are available and include the close relative pair *x*_1_ and *x*_2_ and their mutual relative *y*. The dashed line connecting the MRCA of *x*_1_ and *x*_2_ to *y* indicates the pedigree structure between that MRCA and *y* need not be known. Sexes here are arbitrary and the pedigree relationship type inference works identically for all sample sexes. Haplotypes for the genotyped individuals appear below each pedigree plot as blue or grey vertical bars, with haplotypes for ungenotyped common ancestors of the HS and AV pairs that are related to *y* also shown. The blue regions are either one haplotype of the MRCA of *x*_1_ and *x*_2_, or IBD segments other individuals share with this haplotype. (Grey regions are other genome segments that are not IBD with the blue haplotype and we do not consider them.) The black box outlines the regions shared IBD between *x*_1_ and *y*, and the red box outlines the regions *x*_2_ and *y* share IBD.

In practice, the above ratios vary around their expectations, including for distinct samples *y*. This variability arises both because of the variance in IBD sharing between the close relative pair (i.e., depending on the outcome of the small number of meioses that separate them), and also the variance due to the meioses that separate *y* from the MRCA of *x*_1_ and *x*_2_, with the latter variance increasing for greater meiotic distance. More specifically, mutual relatives *y* with a large meiotic separation will share a relatively small fraction of their genome IBD with the MRCA of *x*_1_ and *x*_2_, with a higher coefficient of variation for this sharing rate than those with lower meiotic distance ^23^, leading to higher variance in the ratios. Therefore, the more closely related *y* is to the MRCA of *x*_1_ and *x*_2_, the more precise the ratios will be.

In large samples, data for multiple mutual relatives *y* can be common, and considering only a single *y* will typically provide less information than combining data from multiple samples. In particular, combining IBD regions from multiple *y* individuals that are related to the same MRCA of *x*_1_ and *x*_2_ will result in a larger fraction of the genome that that MRCA shares IBD to some relative compared to using only one *y*. Our approach to incorporating multiple *y* samples into the ratios is to take the union over each *y* of the three- and two-way IBD sharing regions that form those ratios. This effectively reconstructs the IBD sharing of one or more ungenotyped relative ^17^ that is more closely related to *x*_1_ and *x*_2_ than any single *y*, thereby reducing the variance of the calculated ratios (Figure 2). These ratios are:

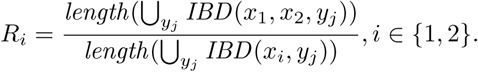

**Figure 2:**
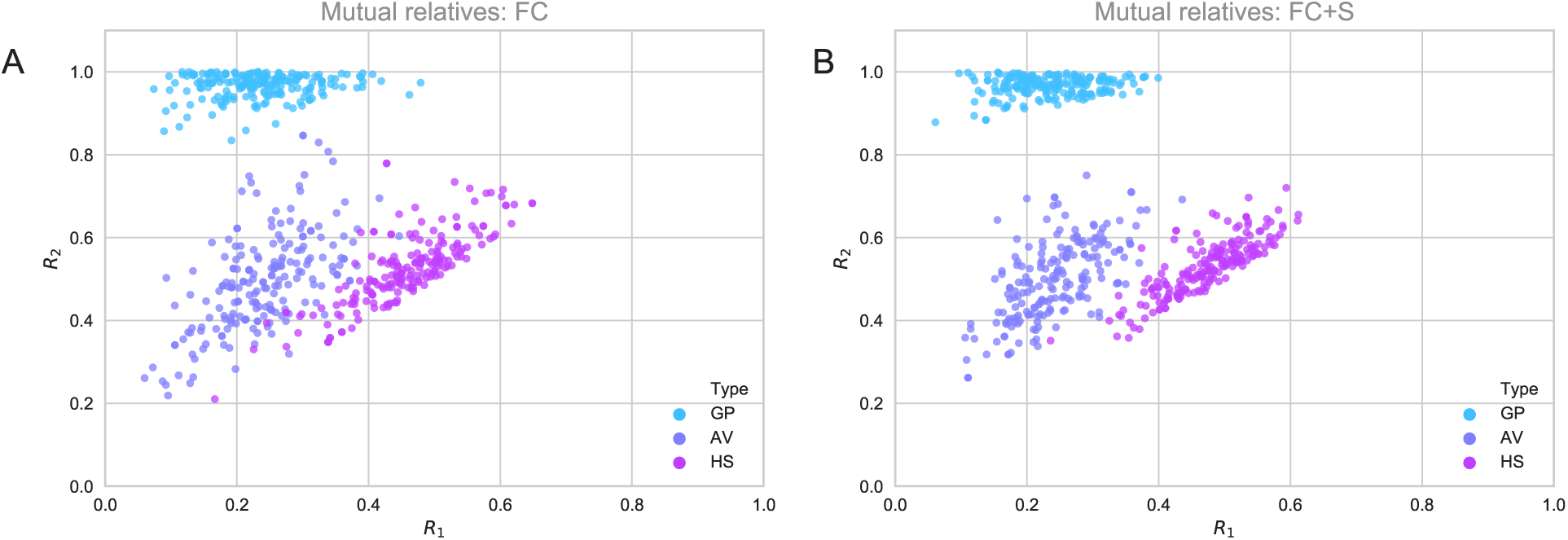
The *R*_1_ and *R*_2_ ratios cluster more tightly when using multiple mutual relatives. Ratios *R*_1_ and *R*_2_ from 200 simulated pairs of each relationship type, calculated using mutual relative(s) of one first cousin (FC) of the genetically older sample in the second degree pair, and (B) combining one first cousin and his/her sibling (FC+S). Here we swap labels if needed so that *R*_1_ *≤ R*_2_.

Here *y*_*j*_ ranges over the mutual relatives that are available in the dataset and satisfy the MRCA assumption. The above assumes that all *y*_*j*_ share IBD segments to only one haplotype of *x*_1_ and/or *x*_2_ at any region, such that the *y*_*j*_ cannot be related through both parents of either *x*_1_ or *x*_2_.

### Classifying relationship types using kernel density estimation models

CREST adopts KDEs to classify the three second degree relationship types using the ratios *R*_1_ and *R*_2_ as features. To train and evaluate the KDEs, for each such relationship type, we first simulated genotype data for a range of pedigree structures that include various mutual relatives, and we derived *R*_1_ and *R*_2_ ratios from the IBD segments that IBIS ^22^ detects in the simulated genotypes (see “Simulations” for details). Because the *R*_1_ and *R*_2_ values are ordered, and since we only seek to classify the relationship types (with directionality considered separately), we exchanged the order of the two ratios such that *R*_1_ ≤ *R*_2_. This shrinks the space the features range over, increasing precision. We then trained separate KDEs for each relationship type and used five-fold cross validation to select their optimal bandwidth and kernel functions.

As noted earlier, the closer the mutual relatives are to the target pair, the less variance the ratios will tend to have, yielding more reliable classification. Therefore, to build models that account for this variance, we incor-porate another feature that is associated with it: what we term the *genome coverage rate, C*, of the pair for a given set of mutual relatives. We define this as 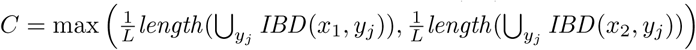, where *L* is the total (M) genetic length of the genome. Thus, it is the larger of either the IBD fraction of *x*_1_ and the mutual relatives or that of *x*_2_. This genome coverage rate is anti-correlated with the variance in the ratios (Figure S1) since it is related to how much of the genome of *x*_1_ and *x*_2_’s MRCA is covered by IBD segments in mutual relatives.

To incorporate genome coverage into our models, we built KDEs stratified by *C*, one for each of several bins. When *C* < 0.2, the bins span intervals of size 0.025, and we use only one bin for *C* ≥ 0.2 because the variances of *R*_1_ and *R*_2_ appear more constant above this threshold (Figure S1). CREST does not attempt to classify pairs with a *C* < 0.025 since distinguishing relationships is difficult with such a low signal. For a given genome coverage bin, we trained KDEs using five-fold cross validation for each bin separately.

To classify a pair’s relationship type, CREST calculates the posterior probability of each type using the likelihood of *R*_1_ and *R*_2_ under the estimated density function for the corresponding type. It outputs these probabilities, calculated as 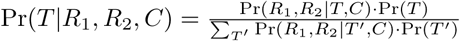, where *T* ∈ {GP, AV, HS} is the type, and Pr(*T*) is the prior probability of the given type, which we take to be 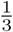 for all *T*. Pr(*R*_1_, *R*_2_|*T, C*) is the likelihood of *R*_1_ and *R*_2_ for a given relationship *T*, and this is from the KDE applicable to the given genome coverage value *C*. As CREST reports all these probabilities, users can choose to use the maximum a posteriori relationship type or to incorporate the probabilities into downstream analyses. In Results, we use the maximum a posteriori type unless otherwise specified.

### Inferring the directionality of the relationship

CREST leverages the ratios *R*_1_ and *R*_2_ to determine the directionality of relationships. More specifically, CREST identifies which sample is the grandparent, aunt/uncle, and parent in GP, AV, and PC pairs, respectively, by comparing their ratios calculated using a set of mutual relatives. In principle, the genetically older sample should inherit more DNA from the MRCA that person shares with each *y*, such that the union of their pairwise IBD sharing over mutual relatives will be greater than that of the younger sample. The latter quantity is in the denominator of the ratios, so CREST determines the directionality as 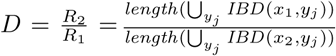. If *D* > 1, *x*_*1*_ should be genetically older than *x*_2_, and the reverse if *D* < 1.

### Inferring the sex of ungenotyped relatives

CREST provides information beyond the relationship type of HS and GP pairs by inferring the sex of ungenotyped relatives: the parents of HS pairs, and the intermediate parent of GP pairs. This inference is possible due to distinct features of male and female maps that lead to different patterns in the IBD segments paternal and maternal pairs share. It is common practice to use a sex-averaged genetic map when analyzing relatives ^13,16,17^, but doing so overlooks the substantial differences between the sex-specific maps. In general, the female map has a greater genetic length (1.6× on the autosomes) than the male map, but the male map is locally longer near the telomeres ^18^. The autosomal length difference between the maps affects the number of crossovers—and thus the number of IBD segments—transmitted through male or female meioses ^24^. Accordingly, the distributions of IBD segment numbers differ meaningfully between paternal and maternal HS and GP relatives (Figure S2), to the extent that limited classification is in principle possible using segment number alone. However, this approach omits the meaningful differences between male and female recombination rates at given physical positions. To best utilize all the information that the sex-specific maps provide, CREST therefore leverages the IBD segment positions and lengths to compute the probability that each segment was transmitted through male or female meioses. In the special cases of HS and GP pairs, the nature of the relationship precludes the possibility that shared IBD segments could have been transmitted by multiple individuals, so this probability naturally infers whether the relationship is paternal or maternal.

By definition, an IBD segment consists of a region of sequence shared between two or more individuals that is flanked by a pair of crossovers, or by the start or end of the chromosome. We make use of the locations of any crossovers implied by IBD segments to infer the transmitting parent’s sex. However, in practice the IBD boundaries will not correspond exactly to the locations of those crossovers ^22,25^. We therefore model the flanking crossovers as falling within windows *w*_0_, *w*_1_ of fixed physical length (Figure 3). This window length is a parameter, but must be small relative to the minimum length of the detected IBD segments. Window size is in some sense a measure of confidence in the IBD segment boundaries; the more accurate the IBD segments, the smaller the window can be, which will increase the precision of the inference (up to the limit of the genetic maps’ resolution).

**Figure 3:**
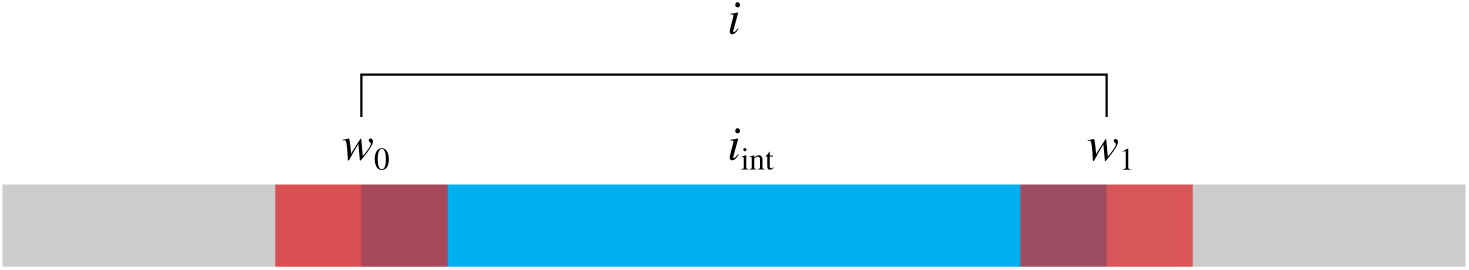
Modeling IBD segments as non-overlapping windows. An IBD segment *i*, in blue, is an interval of sequence shared between two or more individuals. These samples inherit the sequence *i* unrecombined relative to the surrounding non-IBD sequence, in gray. The segment must necessarily be flanked by a pair of crossovers (or by one or both chromosome ends). The precise locations of these crossovers is unknown, because inferred IBD segments have noisy end points. We assume the crossovers fall within some windows *w*_0_ and *w*_1_, in translucent red, centered at the IBD start and end points. The remainder of the segment that is not overlapped by those windows is itself a window we label *i*_int_. Together *w*_0_, *i*_int_ and *w*_1_ approximate *i*.

We model crossing over as a Poisson process for mathematical convenience, although we recognize that this neglects the phenomenon of crossover interference. Recent work provides the means to model interference within arbitrary numbers of meioses from a single genetic map ^24^, but to our knowledge there is no corresponding analytical model capable of handling interference from combinations of both male and female meioses.

As noted throughout, sex detection in CREST is presently applicable only for HS and GP pairs, where the small number of intervening meioses and the presence of only a single intermediate unknown relative simplify modeling considerations. In the case of AV pairs, the IBD segments descend from two grandparents, one male and one female, which complicates the sex inference calculation. More distant relatives pose similar challenges. By contrast, in the case of HS and GP relatives, there is only a single intermediary relative of unknown sex *S* linking the pair: the shared parent for an HS pair, or the parent of the grandchild in a GP pair. That is, the observable recombination events responsible for the flanking crossovers of IBD segments stem only from one relative of unknown sex *S* ∈ {*M, F*}, where *M* is male, *F* is female. Consequently, all crossovers are generated from either a male or female recombination map. For *n* = *n*_*M*_ + *n*_*F*_ meioses separating a HS or GP pair, where *n*_*M*_ and *n*_*F*_ are the number of male and female meioses, respectively, either *n*_*M*_ = 0 or *n*_*F*_ = 0. Consequently *n* = *n*_*S*_, where *S* is the true sex of the intermediate relative we seek to infer.

The genetic length of an interval in Morgans is the number of crossovers expected to occur in that span of sequence during a single meiosis, thus the rate of the crossing over Poisson process in the interval in one meiosis. Suppose a window *w* overlapping part of an IBD segment has a genetic length *C*_*S*_ (*w*) measured on the genetic map of sex *S*. The probability of *k* crossovers occurring within this window over *n*_*S*_ meioses is given by the Poisson mass function with rate *n*_*S*_ ℓ_*S*_ (*w*), i.e.

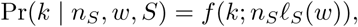

because the rate of the sum of independent Poisson processes is equal to the sum of their rates.

We model IBD segments as a series of three windows, each with known numbers of internal crossovers, that subdivide the interval spanned by an IBD segment *i* into non-overlapping windows (*w*_0_, *i*_int_, *w*_1_) (Figure 3). The windows *w*_0_ and *w*_1_ each contain a single crossover and form the bounds of the IBD segment. In the event that *i* is bounded by either end of a chromosome, CREST omits the corresponding flanking window *w*_0_ or *w*_1_. The remaining interior of *i* falls within *i*_int_. Conveniently, for HS and GP pairs, the IBD segment overlapped by *i*_int_ should contain no crossovers at all in these lineages. This property does not follow for all other relationship types, as internal crossovers can occur in the IBD segments shared by AV pairs, for example. (Here and throughout, we consider IBD segments to continuously span any internal switches between the haplotypes of either individual that carries the segment. This is how most detectors, including IBIS, operate, since distinguishing true haplotype switches from phase switch errors is difficult in population samples.)

The probability of finding one crossover in each window and none in the interior of the interval is:

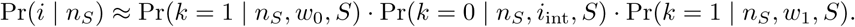

Our prior probabilities that the pair is related maternally or paternally are equal, so 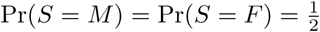. To calculate the joint probability of all IBD segments *I* = {*i*_1_, *i*_2_, …}, we assume that each segment forms independently, which follows from the Poisson model, so Pr(*I* | *n*_*S*_) = Π*i∈I* Pr(*i* | *n*_*S*_).

The above equation gives the probability of observing all IBD segments given some number of meioses *n*_*S*_. However, in the special case of HS and GP pairs, all meioses generating crossovers occur in an individual of sex *S*, i.e., *n* = *n*_*S*_, as noted earlier. Necessarily then all segments must have come from the individual of sex *S*, so Pr(*I* | *S*) = Pr(*I* | *n*_*S*_). This equivalence allows us to apply Bayes’ rule to solve for the probability that the individual is of sex *S*, for *S* ∈ *{M, F}*, given the segments *i* ∈ *I*:

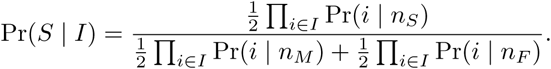

We use this to calculate a logarithm of odds (LOD) score

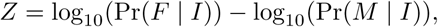

which we use to classify pairs; negative-scoring pairs likely derive from a male, and positive-scoring pairs from a female.

### Simulations

To train and test CREST’s relationship type inference, we simulated a range of pedigree structures that include one GP, AV, or HS pair and one or more of their mutual relatives using Ped-sim ^24^ (Figure S3). In all cases we used sex-specific genetic maps ^18^ and crossover interference ^26^ modeling in these simulations, and a collection of European descent samples ^27^ as the input phased data. (The latter were previously phased using Beagle ^28^, and filtered so that no pair is more closely related than fifth degree ^17^.)

The simulated data we used for training include mutual relatives that vary from first cousins to second cousins of the genetically older sample in the second degree pair. We simulated enough samples to obtain 800 pedigrees within each KDE genome coverage bin. As the coverage rate varies for a given pedigree structure, we simulated 800 pedigrees for each relationship type and pedigree structure class in four batches of 200 pedigrees each. We then mapped these to the corresponding genome coverage bin based on the IBD segments inferred by IBIS, and we randomly downsampled to obtain 800 pedigrees per bin. The pedigrees include nine different combinations of mutual relatives: one first cousin; one first cousin and his/her sibling; two first cousins that also are first cousins to each other; three first cousins that are first cousins to each other; one first cousin and his/her niece/nephew; one first cousin once removed and his/her sibling; one first cousin once removed and his/her niece/nephew; one second cousin; one second cousins and his/her sibling. Thus, we include third degree relatives (first cousins) and as far as seventh degree relatives (second cousins twice removed of a grandchild) for training.

To compare CREST with PADRE ^21^, we also simulated seven different pedigree structures that still include the second degree pairs and with mutual relatives consisting of (again with respect to genetically older sample): one first cousin and his/her sibling (FC+S); one first cousin and his/her child (FC+C); one first cousin and his/her niece/nephew (FC+N); one first cousin once removed and his/her sibling (FC1R+S); one first cousin once removed and his/her child (FC1R+C); one first cousin once removed and his/her niece/nephew (FC1R+N); and one second cousin and his/her sibling (SC+S). We tested both methods using 200 pedigrees of each structure for all three types of second degree relatives.

We further evaluated CREST’s inference accuracy across genome coverage bins. For this analysis, we simulated 200 copies for each relationship type of the same nine pedigree structures we used for training (above). We then mapped these to genome coverage bins and downsampled to obtain 200 copies per bin.

To test CREST’s ability to perform sex inference of the ungenotyped parent linking HS and GP pairs, we simulated 1,800 HS pairs and 900 GP pedigrees, the latter including data from a grandmother-grandfather couple and their grandchild, for a total of 1,800 pairs for each relationship type. We used a feature of Ped-sim to constrain the reproducing sex in each pedigree, such that it generated 900 each of maternal and paternal HS pairs, and 900 maternal and paternal GP pairs (with equal numbers of grandmothers and grandfathers). To increase the sample size, we performed four replicates of these simulations, for a total of 7,200 HS and GP pairs, split evenly into 3,600 maternal and paternal pairs.

### Parameters used to run each method

To collect IBD segments for relationship type inference, we first ran IBIS ^22^ with a minimum segment length of 7 cM, a window size of 0.5 cM, an error threshold value of 0.004, and a marker density of 50 SNPs per cM. Since PADRE requires results from ERSA ^13^ and PRIMUS ^10^ as inputs, we ran them separately on the simulation data. To run PRIMUS (v1.9.0), we first used the `--no_IMUS` and `--no_PR` options, which corresponds to only running PLINK ^29^ (v1.90b2k) to calculate relatedness estimates. We then filtered the output file from PLINK to only include pairs from the same pedigree. Next we ran PRIMUS on this file to infer pedigrees, allowing it to search for up to second degree relatives using the `--degree_rel_cutoff 2` option (all simulation pedigrees it applies to include only first and second degree relatives). Meanwhile, ERSA needs inferred IBD segments from GERMLINE ^30^ as input, while GERMLINE works on phased data, so we also ran Eagle ^31^ v2.4 to phase the simulated genotypes. Each of the Ped-sim simulation runs we used for the PADRE comparison generated data for 200 pedigrees for all three relationship types, and each pedigree includes data from four samples, for a total of 2,400 samples output by Ped-sim. We ran Eagle separately on all seven of these simulated genotype datasets. We then ran GERMLINE on the seven datasets with the options `-err_het 2 -err_hom 1 -min_m 1 -bits 64`. Then we ran ERSA v2.1 with default settings on the GERMLINE output for each dataset.

After all these steps, we ran PADRE v1.0. We found that PADRE initially crashed in some tests, with the source of the crashes being the structure of pedigrees PRIMUS inferred, so we removed the pedigrees that cause the crashes from consideration by PADRE (as in another PADRE analysis ^17^). This avoids calling these tests as PADRE failures, thereby improving its performance.

The runtimes we report are from servers with four Xeon E5 4620 2.20 GHz processors, and we ran IBIS with one thread. (CREST is not multithreaded.)

For sex-inference analyses, we generated IBD segments with IBIS using a minimum segment length of 7 cM, a window size of 0.01 cM, and an error threshold value of 0.04.

### Real data processing

To test CREST’s relative type inference on the GS dataset, we ran IBIS v0.95 (a slightly later version than that used in the simulated data) using default parameters. We noticed some regions of the genome include excess IBD sharing when averaged across all sample pairs, and we used an approach to remove IBD segments that cover these regions from the analysis (below). Following this filter, we inferred degrees of relatedness among pairs (based on the IBD segments) and restricted the analysis to mutual relatives that are third to sixth degree relatives of both members of the target pair. As IBIS is still under development, we further excluded mutual relatives that are inferred as second degree by KING ^14^.

As noted earlier, CREST assumes the mutual relatives do not have IBD sharing to more than one haplotype of the target samples, so CREST first identifies the mutual relative *y*^*^ with the highest kinship coefficient to either member of each pair. It then finds up to fifth degree relatives of *y*^*^ that are also mutual relatives of the target pair and uses all these samples to perform classification.

Because samples can be part of multiple target pairs, we averaged the classification results across all pairs a given individual is part of for each relationship type. We do not average across relationship types, so a given sample can be both a grandchild and a half-sibling with results from the two types considered independent.

To filter IBD segments in regions with excess sharing, we first divided the genome into 1 cM bins and computed the number of IBD segments that overlap each bin. We labeled bins as excessive if they have segment counts *>* 3 standard deviations from the mean. Then, for each IBD segment *i*, we calculated *e*(*i*), the genetic length of the excessive bins it overlaps, and we only retained the segment if *ℓ*(*i*) *-e*(*i*) *>* 7 cM.

The model CREST uses to perform sex inference is very sensitive to errors in the positions of IBD segment breakpoints, and a common issue for IBD segment detection is the inability to detect complete segments ^25^. Detectors frequently identify multiple short segments separated by small gaps rather than one longer contiguous segment. We addressed this issue in two ways. First, we used a relatively lax error threshold parameter in IBIS (0.04, as in the simulations), which should prevent genotyping errors from disrupting a true segment. However, this comes with a trade off, as an overly high error tolerance can detect false segments as well. This is partly remedied by choosing a minimum length of 7 cM for detected segments, since false segments are more likely to be short. The second approach we apply is to post-process segments by “stitching.” Simply put, we stitch, or join, two IBD segments together if they are separated by a sufficiently small gap, which, for the second degree relatives in this analysis, is likely to be erroneous. The maximum stitchable gap we have chosen for the sex inference results in real data was 0.5 cM.

## Results

To evaluate CREST’s ability to distinguish second degree relationship types, we first compared its performance with that of PADRE using simulated pedigrees that include a range of mutual relatives of the second degree pair. We also used simulations to test CREST’s performance across variable genome coverage rates, its ability to infer directionality for PC, AV, and GP pairs, and its potential to classify third degree relationship types, the latter using perfect IBD segment data. Furthermore, we assessed the sex-specific relationship type inference of CREST, initially in simulated data. As we are unaware of another tool to perform sex-specific relationship type inference using autosomal genotypes, we report only CREST’s inference rates for this analysis.

To validate CREST in real data, we analyzed the GS dataset and compared the inferred second degree relatives with the reported relationships. This evaluation includes CREST’s accuracy in inferring sex-specific relationships for the GS samples.

### Classifying second degree relationship types using CREST and PADRE

To compare CREST with PADRE, we tested each method using simulated data from seven different types of mutual relatives defined by their relationship to the genetically older sample: FC+S, FC+C, FC+N, FC1R+S, FC1R+C, FC1R+N, and SC+S (Methods). PADRE infers relatedness between samples in two networks (each containing relatives identified by PRIMUS ^10^) by maximizing the overall composite likelihood, or the product of the likelihood of the pedigree structures PRIMUS infers for the networks and the pairwise relatedness likelihoods for cross-network pairs from ERSA ^13^. As PADRE considers all possible PRIMUS-inferred pedigrees, it analyzes all second degree relationship types that PRIMUS includes in some pedigree, and it considers the PRIMUS pedigree that maximizes the overall composite likelihood to be correct, yielding an inferred second degree relationship type. PRIMUS networks must contain at least two closely related samples, so the mutual relatives are first and second degree relatives of each other. Thus, all the simulated pedigrees we used to compare PADRE and CREST include a network of the target second degree pair and a second network containing two close mutual relatives. However, we note that CREST works even if data is available for only one appropriate mutual relative of the target pair.

We ran both CREST and PADRE on 200 replicates of each of the pedigree structures, and we extracted the relationship PADRE infers for the target second degree pair from its reconstructed pedigrees. As noted in Methods, PADRE crashed for some tests, and we applied a previously used fix ^17^ that enabled it to analyze most of these cases. For the remaining 2.1% of pedigree structures that this fix did not solve, we exclude the case from consideration in PADRE’s results, thus not penalizing its performance. Additionally, CREST does not classify pairs with a coverage rate *C* < 0.025, and we did count these unclassified pairs (0.57% overall) in its performance.

Figure 4 plots the sensitivity and specificity of CREST and PADRE for the seven types of pedigree structures. (Recall that sensitivity is the true positive rate or the fraction of correctly classified pairs for each relationship type, and specificity is one minus the false positive rate.) CREST’s sensitivity (Figure 4A) ranges from 0.953-0.995 for GP, 0.697-0.970 in AV, and 0.785-0.980 in HS pairs across the seven types of mutual relatives. In contrast, PADRE’s sensitivity is 0.399-0.779 for GP, 0.617-0.920 in AV, and 0.734-0.969 in HS pairs. This corresponds to an increase in sensitivity of 0.093-0.237 in CREST across the mutual relative types, averaged over the three target relationships. Turning to specificity (Figure 4B), CREST’s performance rates are 0.954-1.00 in GP, 0.881-0.988 in AV, and 0.881-0.988 in HS compared to PADRE’s rates of 0.954-0.987 in GP, 0.599-0.872 in AV, and 0.812-0.990 in HS. Averaged over the three relationship types, CREST’s specificity is 0.05-0.118 greater than PADRE.

**Figure 4:**
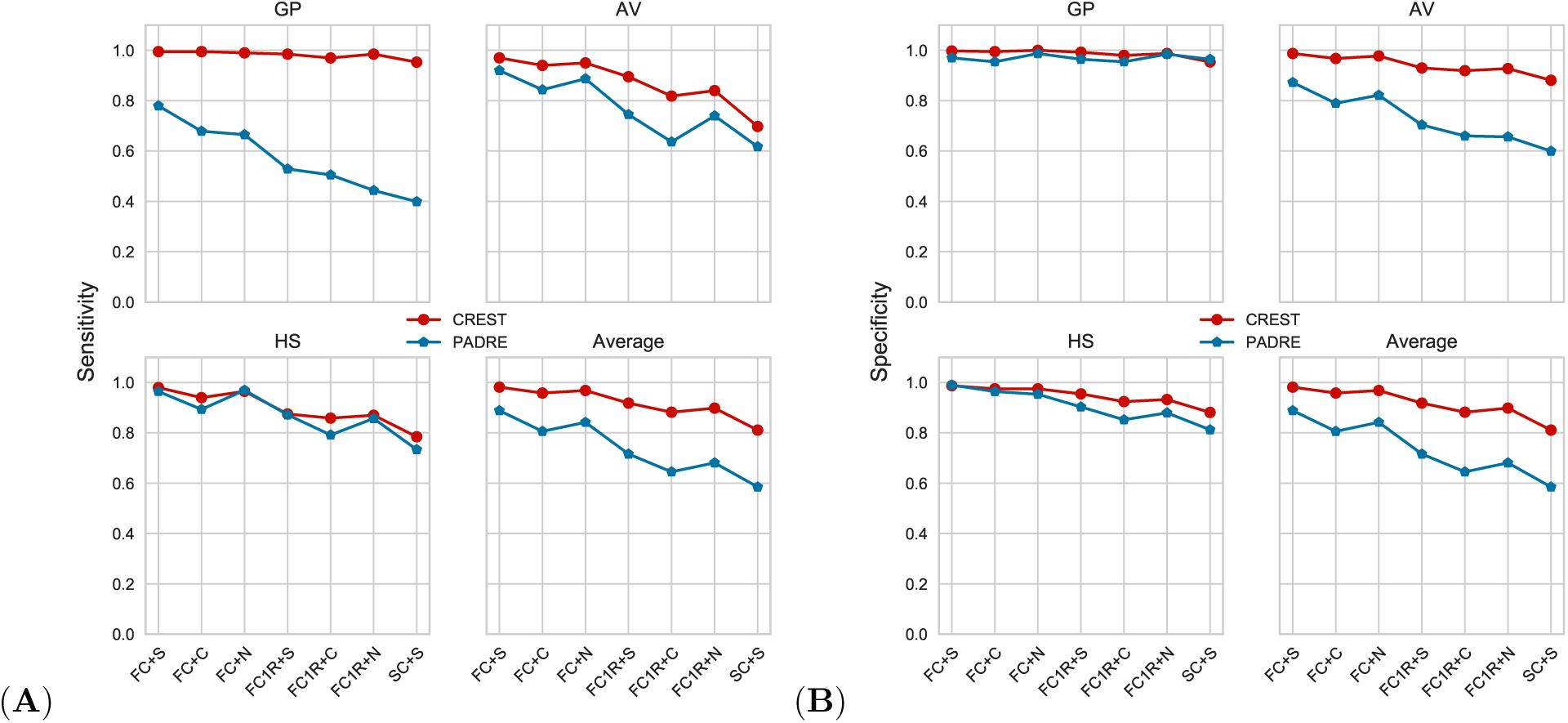
Performance of CREST and PADRE for second degree relationship type classification. (A) The sensitivity and (B) specificity of CREST and PADRE for inferring GP, AV, and HS relationship types, along with the average of these rates across the three relationships. The x-axis indicates the mutual relatives types included in the analysis, with each target relationship type and mutual relative combination including data from 200 pairs. The average of sensitivity and specificity are calculated are across all three relationship types. Analyses that crashed in PADRE are excluded from PADRE’s performance, and those with *C* < 0.025 are excluded from CREST’s.

These results indicate that, for the types of mutual relatives we tested, CREST has higher sensitivity than PADRE for identifying AV and GP pairs. For HS pairs, both CREST and PADRE have good performance, which may be because the mutual relatives are closer to both the target samples in this case compared to other two types. Alternatively, previous work indicated that PADRE may be biased against GP relationship classification and in favor of HS ^17^. Related to this, the confusion matrices show that PADRE misclassified more GP as AV for more distant mutual relatives (Figure S4). In turn, CREST tends to mix HS and AV classification, and is better at identifying GP pairs.

Considering the runtime of these analyses, the IBD detector IBIS ran on one of the seven sets of 2,400 samples (simulated for each of the mutual relative type classes; Methods) in 35.6 CPU minutes, and CREST completed the classification of the relationships in another 5.5 minutes. However, PADRE requires that the samples be phased, have IBD detected (with GERMLINE), and analyzed using both PRIMUS and ERSA. Phasing and ERSA together take more than two CPU days to finish for one of the pedigree types.

### The performance of CREST under variable genome coverage rates

As discussed in Methods and depicted in Figure 4, classification using close mutual relatives has better performance than using more distant relatives. To ensure that the KDE distributions more accurately represent the true relationship probabilities for a given target pair and their mutual relatives, we trained stratified KDEs based on the genome coverage rate *C* of a set of mutual relatives (Methods).

Figure 5 shows the sensitivity and specificity of CREST across the bins of genome coverage rates on which we trained separate KDEs. As expected, the sensitivity and specificity both increase as the coverage grows. For coverage rates between 0.125-0.15, or roughly that expected for one first cousin, CREST’s sensitivity and specificity are both 1.00 for GP, 0.94 and 0.95 for AV, and 0.91 and 0.97 in HS pairs, respectively. Even when *C* is in the lowest bin of 0.025-0.05, CREST still achieves a sensitivity of 0.970 and specificity of 0.960 for GP, 0.670 sensitivity and 0.690 specificity in AV, and 0.710 sensitivity and 0.700 specificity in HS pairs. Notably, the inference of GP pairs generally has quite high sensitivity and specificity regardless of the genome coverage rate. This is likely because, if *x*_*i*_ is the grandchild, *R*_*i*_ = 1, with no variance from the meioses that separate *x*_*i*_ from the grandparent, but only due to false positive and/or false negative IBD segments.

**Figure 5:**
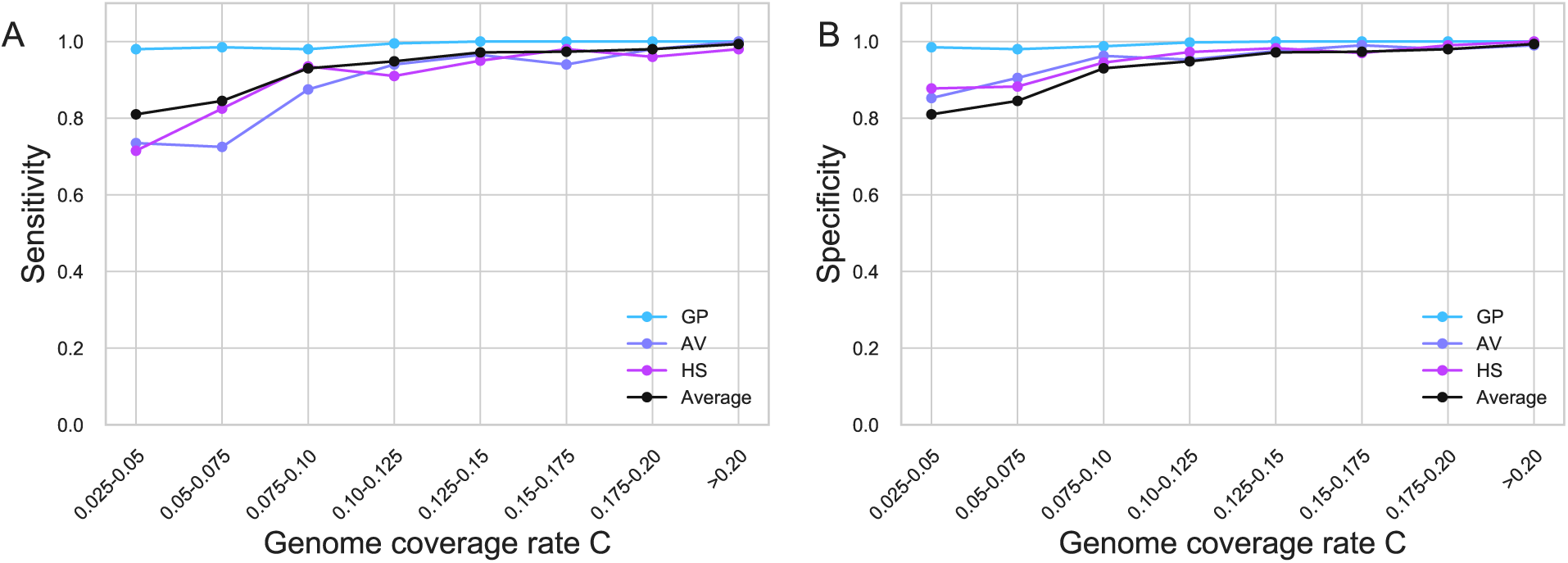
CREST performance across bins of genome coverage rates. (A) The sensitivity and (B) specificity within genome coverage rate (*C*) bins for GP, AV, and HS pairs, and the average across these three types.

The results above consider only the highest posterior probability relationship for the performance evaluations, but this probability is informative about CREST’s confidence in its classification. We tested the accuracy of CREST by only classifying those pairs with posterior probability above a given threshold, omitting pairs below this number from consideration. Figure S5 depicts the average across the three relationship types of F1 scores with the given threshold varying from 0.50 to 0.90. The results show that for low *C* values, increasing the probability threshold is effective at improving the F1 score.

### Detecting the directionality of relationships

To test CREST’s ability to detect the directionality of relationships, we used the same simulated data as in the above genome coverage analysis, but instead of analyzing HS pairs, we used the parent and one member of each HS pair to analyze the performance of PC directionality inference. This analysis includes 200 pairs of GP, AV, and PC relatives, which we analyzed across the same genome coverage rate bins as define CREST’s KDEs. We calculated the *R*_1_ and *R*_2_ values for each pair and used these to infer which sample is the grandparent, aunt/uncle, or parent (Methods). As shown in Figure S6, when *C >* 0.025, which corresponds roughly to using one fifth degree or more informative mutual relative, CREST achieved sensitivity of 1.00 in determining the directionality of GP pairs, 0.99 for AV, and 1.00 for PC pairs.

### CREST has the potential to infer third degree relationship types

In principle, the CREST approach need not be limited to second degree relationships, as a similar logic applies to more distant relatives. To analyze the potential for CREST to distinguish third degree relatives, we tested its ability to classify four third degree relationship types: great-grandparent (GGP), grand-avuncular (GAV), half-avuncular (HAV), and first cousin (FC). Assuming that *x*_1_ is the genetically older sample, for a GGP pair, 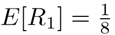 and *E*[*R*_2_] = 1; for a GAV pair, 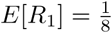 and 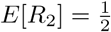; for a HAV pair, 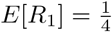 and 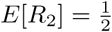; and for a FC pair, 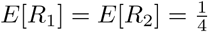.

To conduct this analysis, we employed perfect IBD segments simulated by Ped-sim, and we only included those *>* 5 cM long, envisioning the potential for obtaining extremely reliable IBD segments above this threshold. To train and test this extension of CREST, we simulated 1,000 pedigrees for each of the third degree relationships, with each pedigree including two first cousins of genetically older sample as mutual relatives. After calculating *R*_1_ and *R*_2_, we trained the KDEs using 800 samples and five-fold cross validation for each type. We then tested on the remaining 200 pairs, and found that the inference accuracy is high. The sensitivities are 1.00 for GGP, 0.943 for GAV, 0.932 for HAV, 0.988 for FC pairs (Figure S7). Thus, given sufficient mutual relative data and, in this case, exact IBD segments, CREST can infer third degree relationship types. This is true even when limiting to segments *>* 5 cM long, and may even now find utility when sequence data are available for sets of close relatives that include third degree pairs.

### Sex-specific classification

To evaluate CREST’s sex-specific classification, we used Ped-sim to generate 3,600 maternal and 3,600 paternal pairs for each of the HS and GP relationship types (Methods). For inference, we chose 500 kb as the size of the windows in which we model crossover locations (Figure 3). Figure 6A plots the resulting LOD scores (*Z*), which are positive if the pair is inferred as maternal, and negative for paternal, with a greater magnitude of *Z* corresponding to greater confidence in the call. For the HS pairs, CREST correctly infers the sex of the parent in 99.1% (*N* = 3569) of maternal pairs and 98.0% (*N* = 3528) of paternal pairs. For the GP pairs, CREST infers 74.6% (*N* = 2658) of maternal pairs and 99.4% (*N* = 3578) of paternal pairs correctly. We used the LOD scores to generate receiver operating characteristics (ROCs) for these relationship types, as shown in Figure 6B. The area under the curve (AUC) of these ROCs are high, with an AUC for HS pairs of 0.999 and for GP of 0.986, consistent with the classification results highlighted above.

**Figure 6:**
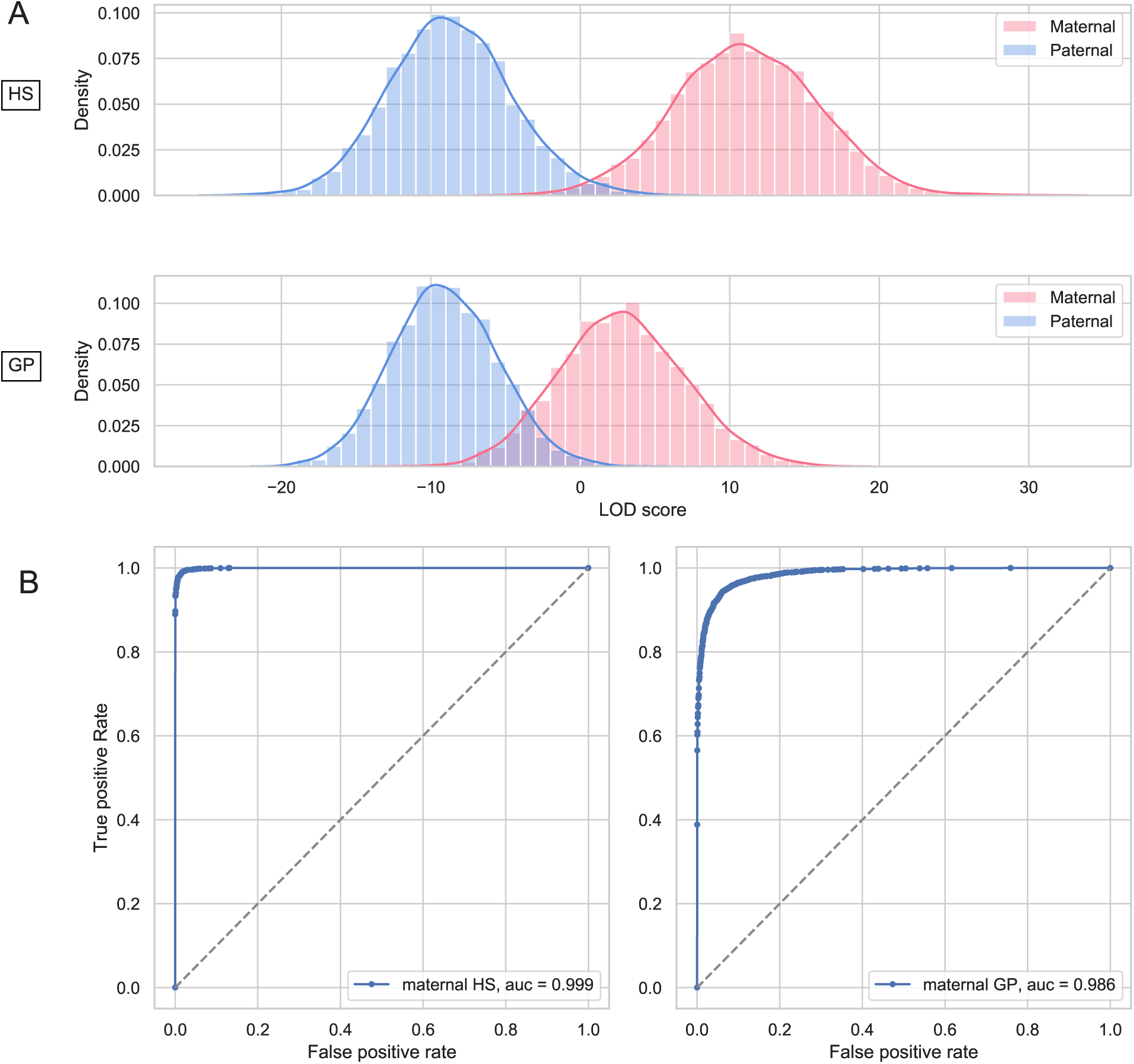
Performance of the sex relationship type inference in simulated data. (A) Histograms showing the distributions of LOD scores for the inferred parental sex of HS (top) and GP (bottom) pairs. Pink samples represent scores for pairs where the true sex is female, and likewise blue for male. (B) ROC curves for HS (left) and GP (right) pairs. The area under the curve (AUC) is provided in the label.

### Validation in Generation Scotland data

In order to test our model in real data, we used CREST to classify second degree relationships in the GS samples, which are enriched in close relatives and include pedigree structures. Considering those pairs with at least one appropriate mutual relative (Methods), this analysis considered 205 GP, 1,949 AV, and 121 HS pairs. For this, we excluded the two HS pairs that the below sex inference analysis flagged as having an uncertain relationship. We applied our KDEs (trained using simulated data) to infer the types of these second degree pairs. Performing this analysis took 8.7 CPU hours to run IBIS, 1.4 CPU hours to filter the IBD segments and put them in the desired format (Methods), and 14.0 CPU minutes to run CREST.

Figures 7A and 7B plot the sensitivity and specificity of this analysis across different genome coverage rates *C*. As expected, both the sensitivity and specificity tend to increase with *C*. However, the sensitivity of GP pairs in the *C >* 0.15 bin drops relative to the next lower coverage bin, and is not as high as in the simulations. This bin has only six true GP pairs with one (effective) misclassified pair (after averaging); the misclassification is due to CREST using a mutual relative that is another grandchild and inferred as a third degree relative of the grandparent. Overall, for *C >* 0.10, CREST’s sensitivity is relatively high with a value of 0.990 for GP, 0.857 for AV, and 0.950 for HS pairs. Similarly, the specificity is high in this coverage range, with values of 0.980 for GP, 0.970 for AV, and 0.890 for HS pairs.

**Figure 7:**
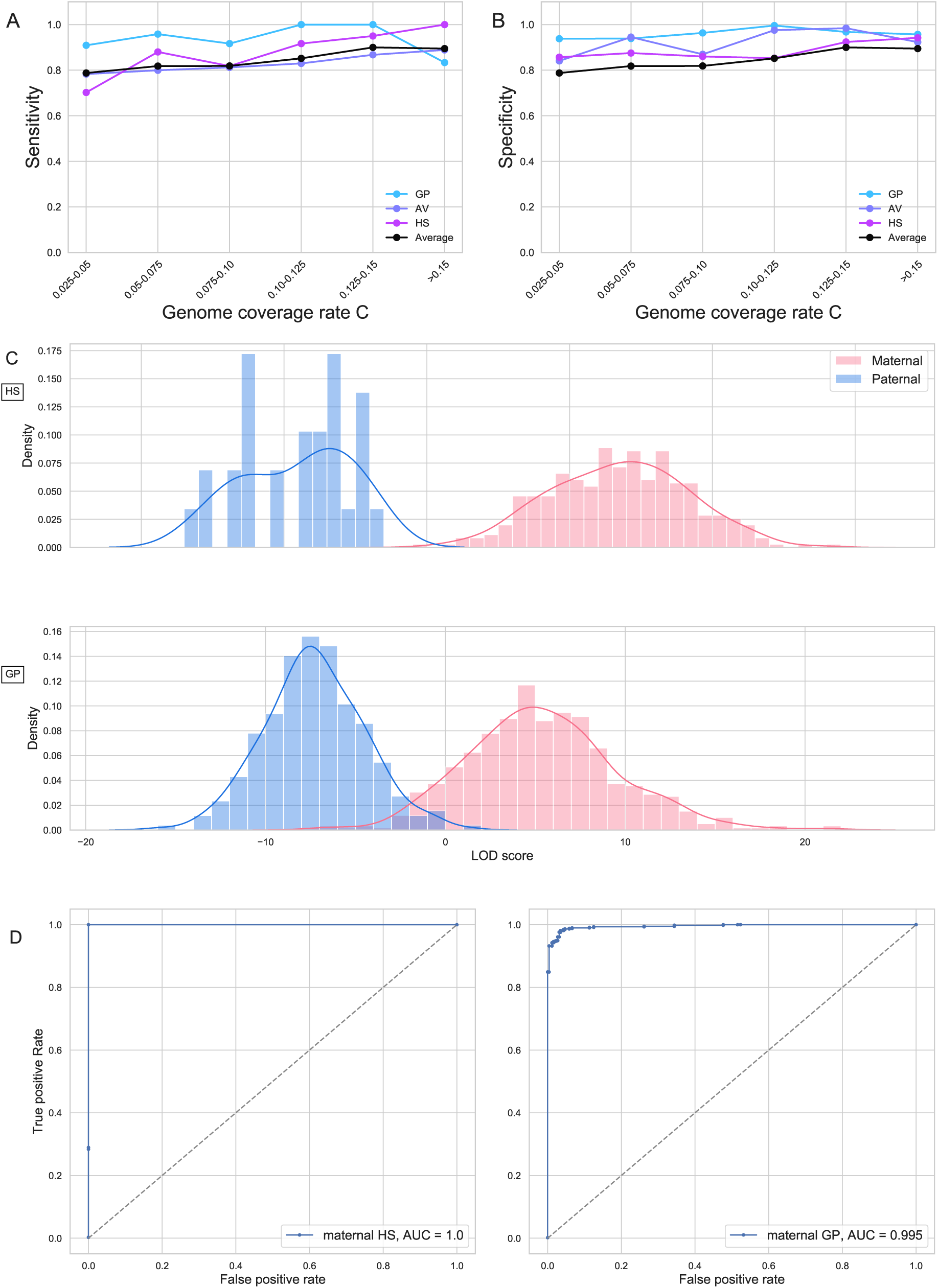
CREST performance on Generation Scotland Data. (A) The sensitivity and (B) specificity of relationship type classification for GP, AV, HS, and the average across these three types in the GS dataset. (C) Histograms showing the distributions of LOD scores for the inferred parental sex of HS (top) and GP (bottom) pairs. Pink samples represent scores for pairs where the true sex is female, and likewise blue for male. (D) ROC curves for HS (left) and GP (right) pairs. The area under the curve (AUC) is provided in the label.

We also used the GS pedigrees to validate the capacity of CREST’s maternal and paternal relationship inference in real data. All 848 GP and 381 HS pairs in the dataset are suitable for this analysis, and we again used flanking windows of 500 kb for inference. After performing this analysis, we noticed that five of the pairs (four HS and one GP pair) had extremely anomalous scores in conflict with their recorded relationships (Figure S8). Of these, we found that two HS pairs were incorrectly reported, and the GP pair was incorrectly listed as such when it is in fact AV. We further concluded that the other two HS pairs are likely of some other relationship type.

After removing the non-HS and GP pairs and updating the mislabeled HS pairs, the LOD scores generally reflect the true relationship types (Figure 7C). Moreover, CREST correctly infers 99.7% (*N* = 348) of maternal HS pairs, 100% (*N* = 29) of paternal HS pairs, 90.8% (*N* = 536) of maternal GP pairs, and 99.6% (*N* = 255) of paternal GP pairs. These results are meaningfully better than those in simulated data, which could be due inaccuracies of the simulation model, including Ped-sim’s genotyping error model. The ROCs for these GS results also reflect the effectiveness of the LOD scores, with AUCs for the HS pairs of 1.00, and 0.995 for the GP pairs (Figure 7D).

## Discussion

Pedigrees have wide ranging utility throughout genetics, with modeling of the transmission of haplotypes among samples central to both linkage analysis and very recent heritability estimation procedures ^6,7^. Data from families are also needed to identify *de novo* recombinations ^18,26,32^ and mutations ^33,34^, and to enable family-based phasing and imputation, the gold-standard means of addressing these problems ^35^.

Given these applications, several methods exist for pedigree reconstruction, and for confirming or disproving reported pedigree relationships ^10–12,16,21,36^. However, differentiating among the relationships that map to a given degree of relatedness has remained challenging. Pairwise relatedness measures, the standard signal for detecting relatives until very recently ^15^, have limited information to enable the classification of relationship types ^16^.

We developed CREST, an approach that infers both pedigree relationships and whether HS and GS pairs are maternally or paternally connected. This latter inference relies on sex-specific genetic maps ^18^, whose genome-wide rate differences were observed in early human genetics analyses ^37^. The male and female maps also differ markedly in their local crossover rates, with these differences forming a key basis to the signals CREST uses for its inference. For example, IBD segment counts, a quantity affected by genome-wide crossover rates, overlap meaningfully in paternal and maternal GP pairs (Figure S2), whereas the LOD scores we calculate show greater separation (Figures 6, 7). We are unaware of prior work that uses sex-specific maps to infer familial relationships in this manner.

CREST’s relationship type inference assumes that mutual relatives connect to a target relative pair through the MRCA of the target pair. To enforce this assumption, which is most readily violated by descendants of the target pair, CREST does not analyze first and second degree relatives of the target pair. However, such close relatives carry IBD segments that span a large fraction of the genome, and so have the potential to be very informative for this inference. We view the inclusion of such samples as of interest for future work.

As genetic testing companies remain popular and provide customers with estimated relationships among samples, CREST has utility in several ways. Most apparently, it can enable the companies to report specific relationship types, including which parent an individual is related through for the relationships CREST applies to. Additionally, while the mutual relatives of a target pair inform the pedigree structure between the pair, providing this pedigree structure to the method DRUID enables more exact detection of the distance between those close relatives and the mutual relatives ^17^. Thus, an iterative procedure is possible, with mutual relatives of unknown relationship to a set of close relatives enabling the detection of the latter pairs’ relationship types, and the resulting pedigrees enabling more precise characterization of their distance to the mutual relatives.

Lastly, a key factor influencing CREST’s performance is the genome coverage rate of the available mutual relatives. More closely related pairs generally have a higher genome coverage. In consequence, with ever increasing sample sizes—and therefore datasets with greater numbers of relatives in general and more close relatives—CREST will infer relationship types with greater reliability going forward.

## Acknowledgements

We thank Archie Campbell for help in evaluating the relationships of Generation Scotland individuals, Daniel Seidman for support in using IBIS, Giulio Genovese for his observations about inferring parent-child directionality, and Shai Carmi for discussions regarding time-dependent Poisson rates. Funding for this work was from an Alfred P. Sloan Research Fellowship and a seed grant from Nancy and Peter Meinig to A.L.W. C.H. is supported by an MRC University Unit Programme Grant MC UU 00007/10 (QTL in Health and Disease). Computing was performed on a cluster administered by the Biotechnology Resource Center at Cornell University. Generation Scotland received core support from the Chief Scientist Office of the Scottish Government Health Directorates [CZD/16/6] and the Scottish Funding Council [HR03006]. Genotyping of the GS:SFHS samples was carried out by the Genetics Core Laboratory at the Edinburgh Clinical Research Facility, University of Edinburgh, Scotland and was funded by the Medical Research Council UK and the Wellcome Trust (Wellcome Trust Strategic Award “STratifying Resilience and Depression Longitudinally” (STRADL) Reference 104036/Z/14/Z). This study makes use of data generated by the Wellcome Trust Case-Control Consortium. A full list of the investigators who contributed to the generation of the data is available from www.wtccc.org.uk. Funding for the project was provided by the Wellcome Trust under award 076113, 085475 and 090355.

## References

[1] Clare Bycroft, Colin Freeman, Desislava Petkova, Gavin Band, Lloyd T. Elliott, Kevin Sharp, Allan Motyer, Damjan Vukcevic, Olivier Delaneau, Jared O’Connell, Adrian Cortes, Samantha Welsh, Alan Young, Mark Effingham, Gil McVean, Stephen Leslie, Naomi Allen, Peter Donnelly, and Jonathan Marchini. The UK Biobank resource with deep phenotyping and genomic data. Nature, 562(7726):203–209, 2018.

[2] Jeffrey Staples, Evan K. Maxwell, Nehal Gosalia, Claudia Gonzaga-Jauregui, Christopher Snyder, Alicia Hawes, John Penn, Ricardo Ulloa, Xiaodong Bai, Alexander E. Lopez, Cristopher V. Van Hout, Colm O’Dushlaine, Tanya M. Teslovich, Shane E. McCarthy, Suganthi Balasubramanian, H. Lester Kirchner, Joseph B. Leader, Michael F. Murray, David H. Ledbetter, Alan R. Shuldiner, George D. Yancoupolos, Frederick E. Dewey, David J. Carey, John D. Overton, Aris Baras, Lukas Habegger, and Jeffrey G. Reid. Profiling and leveraging relatedness in a precision medicine cohort of 92,455 exomes. The American Journal of Human Genetics, 102(5):874–889, May 2018.

[3] Benjamin F Voight and Jonathan K Pritchard. Confounding from cryptic relatedness in case-control association studies. PLOS Genetics, 1(3):e32, 2005.

[4] Hyun Min Kang, Jae Hoon Sul, Susan K. Service, Noah A. Zaitlen, Sit-yee Kong, Nelson B. Freimer, Chiara Sabatti, and Eleazar Eskin. Variance component model to account for sample structure in genome-wide association studies. Nat Genet, 42(4):348–354, Apr 2010.

[5] Xiang Zhou and Matthew Stephens. Genome-wide efficient mixed-model analysis for association studies. Nat Genet, 44(7):821–824, Jul 2012.

[6] Noah Zaitlen, Peter Kraft, Nick Patterson, Bogdan Pasaniuc, Gaurav Bhatia, Samuela Pollack, and Alkes L. Price. Using extended genealogy to estimate components of heritability for 23 quantitative and dichotomous traits. PLOS Genetics, 9(5):e1003520. 05 2013.

[7] Alexander I. Young, Michael L. Frigge, Daniel F. Gudbjartsson, Gudmar Thorleifsson, Gyda Bjornsdottir, Patrick Sulem, Gisli Masson, Unnur Thorsteinsdottir, Kari Stefansson, and Augustine Kong. Relatedness disequilibrium regression estimates heritability without environmental bias. Nature Genetics, 50(9):1304–1310, 2018.

[8] John Wakeley, Léandra King, Bobbi S Low, and Sohini Ramachandran. Gene genealogies within a fixed pedigree, and the robustness of kingman’s coalescent. Genetics, 190(4):1433–1445, 2012.

[9] Elizabeth A Thompson. Identity by descent: variation in meiosis, across genomes, and in populations. Genetics, 194(2):301–326, 2013.

[10] Jeffrey Staples, Dandi Qiao, Michael H. Cho, Edwin K. Silverman, Deborah A. Nickerson, and Jennifer E. Below. Primus: Rapid reconstruction of pedigrees from genome-wide estimates of identity by descent. The American Journal of Human Genetics, 95(5):553–564, Nov 2014.

[11] Amy Ko and Rasmus Nielsen. Composite likelihood method for inferring local pedigrees. PLOS Genetics, 13(8):e1006963. 08 2017.

[12] D. He, Z. Wang, L. Parida, and E. Eskin. Iped2: Inheritance path based pedigree reconstruction algorithm for complicated pedigrees. IEEE/ACM Transactions on Computational Biology and Bioinformatics, 14(5):1094–1103, Sep. 2017.

[13] Chad D. Huff, David J. Witherspoon, Tatum S. Simonson, Jinchuan Xing, W. Scott Watkins, Yuhua Zhang, Therese M. Tuohy, Deborah W. Neklason, Randall W. Burt, Stephen L. Guthery, Scott R. Woodward, and Lynn B. Jorde. Maximum-likelihood estimation of recent shared ancestry (ersa). Genome Research, 21(5):768–774, 2011.

[14] Ani Manichaikul, Josyf C. Mychaleckyj, Stephen S. Rich, Kathy Daly, Michéle Sale, and Wei-Min Chen. Robust relationship inference in genome-wide association studies. Bioinformatics, 26(22):2867–2873, 2010.

[15] Monica D. Ramstetter, Thomas D. Dyer, Donna M. Lehman, Joanne E. Curran, Ravindranath Duggirala, John Blangero, Jason G. Mezey, and Amy L. Williams. Benchmarking relatedness inference methods with genome-wide data from thousands of relatives. Genetics, 207(1):75–82, 2017.

[16] Michael P. Epstein, William L. Duren, and Michael Boehnke. Improved inference of relationship for pairs of individuals. The American Journal of Human Genetics, 67(5):1219–1231, Nov 2000.

[17] Monica D. Ramstetter, Sushila A. Shenoy, Thomas D. Dyer, Donna M. Lehman, Joanne E. Curran, Ravindranath Duggirala, John Blangero, Jason G. Mezey, and Amy L. Williams. Inferring identical-by-descent sharing of sample ancestors promotes high-resolution relative detection. The American Journal of Human Genetics, 103(1):30–44, 2018.

[18] Claude Bhérer, Christopher L Campbell, and Adam Auton. Refined genetic maps reveal sexual dimorphism in human meiotic recombination at multiple scales. Nature Communications, 8, 2017.

[19] Blair H Smith, Archie Campbell, Pamela Linksted, Bridie Fitzpatrick, Cathy Jackson, Shona M Kerr, Ian J Deary, Donald J MacIntyre, Harry Campbell, Mark McGilchrist, Lynne J Hocking, Lucy Wisely, Ian Ford, Robert S Lindsay, Robin Morton, Colin N A Palmer, Anna F Dominiczak, David J Porteous, and Andrew D Morris. Cohort profile: Generation Scotland: Scottish family health study (GS:SFHS). The study, its participants and their potential for genetic research on health and illness. International Journal of Epidemiology, 42(3):689–700, 2013.

[20] Reka Nagy, Thibaud S. Boutin, Jonathan Marten, Jennifer E. Huffman, Shona M. Kerr, Archie Campbell, Louise Evenden, Jude Gibson, Carmen Amador, David M. Howard, Pau Navarro, Andrew Morris, Ian J. Deary, Lynne J. Hocking, Sandosh Padmanabhan, Blair H. Smith, Peter Joshi, James F. Wilson, Nicholas D. Hastie, Alan F. Wright, Andrew M. McIntosh, David J. Porteous, Chris S. Haley, Veronique Vitart, and Caroline Hayward. Exploration of haplotype research consortium imputation for genomewide association studies in 20,032 Generation Scotland participants. Genome Medicine, 9(1):23, Mar 2017.

[21] Jeffrey Staples, David J Witherspoon, Lynn B Jorde, Deborah A Nickerson, Jennifer E Below, Chad D Huff, University of Washington Center for Mendelian Genomics, et al. PADRE: Pedigree-aware distant-relationship estimation. The American Journal of Human Genetics, 99(1):154–162, 2016.

[22] Daniel N. Seidman, Sushila A. Shenoy, Minsoo Kim, Ramya Babu, Thomas D. Dyer, Donna M. Lehman, Joanne E. Curran, Ravindranath Duggirala, and John Blangero Amy L. Williams. Rapid, phase-free detection of long identical by descent segments enables fast relationship classification. (Under review), 2019.

[23] WG Hill and BS Weir. Variation in actual relationship as a consequence of Mendelian sampling and linkage. Genetics Research, 93(01):47–64, 2011.

[24] Madison Caballero, Daniel N. Seidman, Jens Sannerud, Thomas D. Dyer, Donna M. Lehman, Joanne E. Curran, Ravindranath Duggirala, John Blangero, Shai Carmi, and Amy L. Williams. Crossover interference and sex-specific genetic maps shape identical by descent sharing in close relatives. bioRxiv, 2019.

[25] Douglas W. Bjelland, Uday Lingala, Piyush S. Patel, Matt Jones, and Matthew C. Keller. A fast and accurate method for detection of ibd shared haplotypes in genome-wide snp data. European Journal Of Human Genetics, 25:617, Feb 2017. Article.

[26] Christopher L. Campbell, Nicholas A. Furlotte, Nick Eriksson, David Hinds, and Adam Auton. Escape from crossover interference increases with maternal age. Nature Communications, 6:6260, Feb 2015.

[27] International Multiple Sclerosis Genetics Consortium, Wellcome Trust Case Control Consortium 2, et al. Genetic risk and a primary role for cell-mediated immune mechanisms in multiple sclerosis. Nature, 476(7359):214–219, 2011.

[28] Brian L. Browning and Sharon R. Browning. A unified approach to genotype imputation and haplotype-phase inference for large data sets of trios and unrelated individuals. The American Journal of Human Genetics, 84(2):210–223, 2009.

[29] Christopher C. Chang, Carson C. Chow, Laurent CAM Tellier, Shashaank Vattikuti, Shaun M. Purcell, and James J. Lee. Second-generation plink: rising to the challenge of larger and richer datasets. GigaScience, 4(1):7, 2015.

[30] Alexander Gusev, Jennifer K. Lowe, Markus Stoffel, Mark J. Daly, David Altshuler, Jan L. Breslow, Jeffrey M. Friedman, and Itsik Pe’er. Whole population, genome-wide mapping of hidden relatedness. Genome Research, 19(2):318–326, 2009.

[31] Po-Ru Loh, Petr Danecek, Pier Francesco Palamara, Christian Fuchsberger, Yakir A Reshef, Hilary K Finucane, Sebastian Schoenherr, Lukas Forer, Shane McCarthy, Goncalo R. Abecasis, Richard Durbin, and Alkes L Price. Reference-based phasing using the haplotype reference consortium panel. Nat Genet, 48(11):1443–1448, Nov 2016.

[32] Bjarni V. Halldorsson, Gunnar Palsson, Olafur A. Stefansson, Hakon Jonsson, Marteinn T. Hardarson, Hannes P. Eggertsson, Bjarni Gunnarsson, Asmundur Oddsson, Gisli H. Halldorsson, Florian Zink, Sigurjon A. Gudjonsson, Michael L. Frigge, Gudmar Thorleifsson, Asgeir Sigurdsson, Simon N. Stacey, Patrick Sulem, Gisli Masson, Agnar Helgason, Daniel F. Gudbjartsson, Unnur Thorsteinsdottir, and Kari Stefansson. Characterizing mutagenic effects of recombination through a sequence-level genetic map. Science, 363(6425), 2019.

[33] Raheleh Rahbari, Arthur Wuster, Sarah J. Lindsay, Robert J. Hardwick, Ludmil B. Alexandrov, Saeed Al Turki, Anna Dominiczak, Andrew Morris, David Porteous, Blair Smith, Michael R. Stratton, UK 10K Consortium, and Matthew E. Hurles. Timing, rates and spectra of human germline mutation. Nature Genetics, 48:126–133, Dec 2015. Article.

[34] Thomas A. Sasani, Brent S. Pedersen, Ziyue Gao, Lisa Baird, Molly Przeworski, Lynn B. Jorde, and Aaron R. Quinlan. Large, three-generation ceph families reveal post-zygotic mosaicism and variability in germline mutation accumulation. bioRxiv, 2019.

[35] Sharon R Browning and Brian L Browning. Haplotype phasing: existing methods and new developments. Nature Reviews Genetics, 12(10):703–714, 2011.

[36] Lei Sun and Apostolos Dimitromanolakis. PREST-plus identifies pedigree errors and cryptic relatedness in the GAW18 sample using genome-wide SNP data. BMC Proceedings, 8(Suppl 1):S23, 2014.

[37] J. H. Renwick. The mapping of human chromosomes. Annual Review of Genetics, 5(1):81–120, 1971. PMID: 16097652.

