## Supplementary Material for "Distinguishing pedigree relationships using multi-way identical by descent sharing and sex-specific genetic maps"

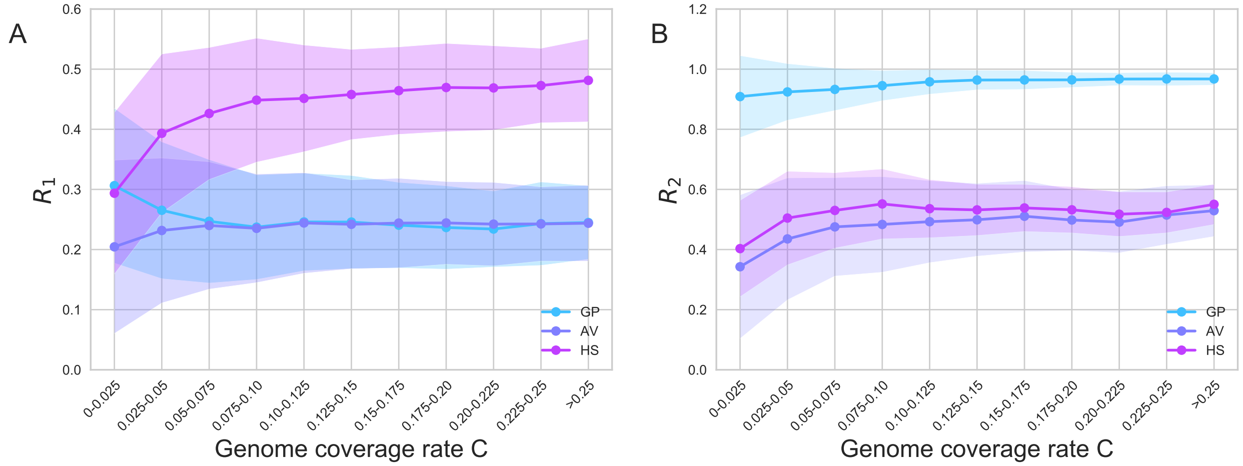


**Figure S1: The variance of ratios $R_1$ and $R_2$ decrease as the genome coverage rate increases.** The dots show the mean ratio values for every bin. The shaded regions are one standard deviation from the mean. Results are from simulated data, with IBD segments detected in genotype data, for all three types.

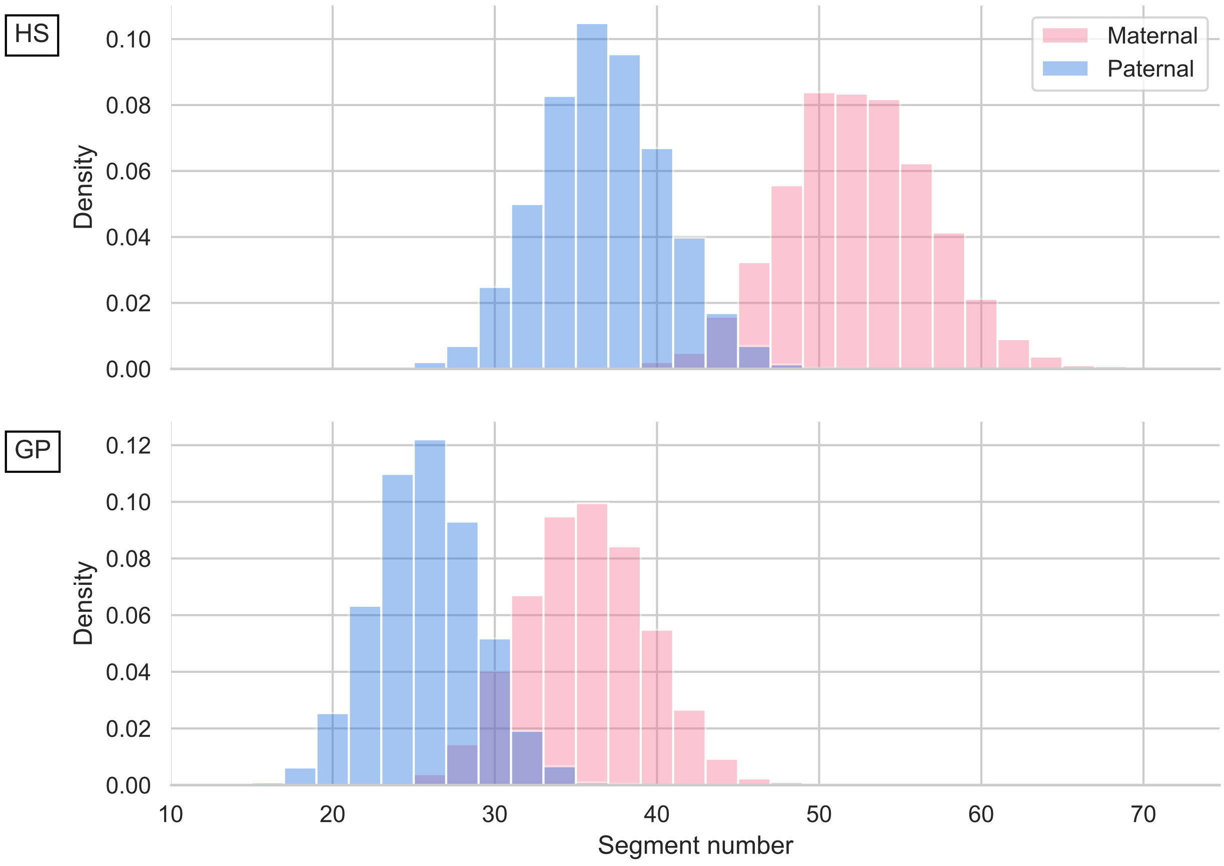


**Figure S2: Segment number distributions for simulated HS and GP pairs.** Histograms of total IBD segment numbers between HS (top) and GP (bottom) pairs using IBD segments generated by Ped-sim.

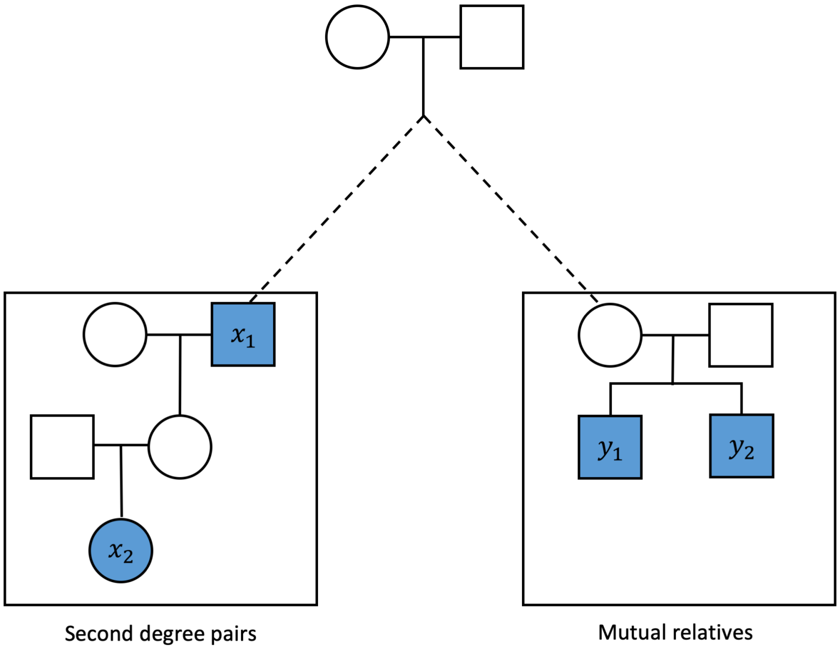


**Figure S3: The structure of the simulated pedigrees used to evaluate CREST's relationship type classification.** The left side is the second degree pair, which is either a GP, AV, or HS pair. The right side are the mutual relatives which include one or more individuals with relationships to the second degree pair and to each other described in Methods. Genotyped samples are shown as filled shapes. The dashed line connects the second degree pair and the mutual relatives to their unknown MRCA.

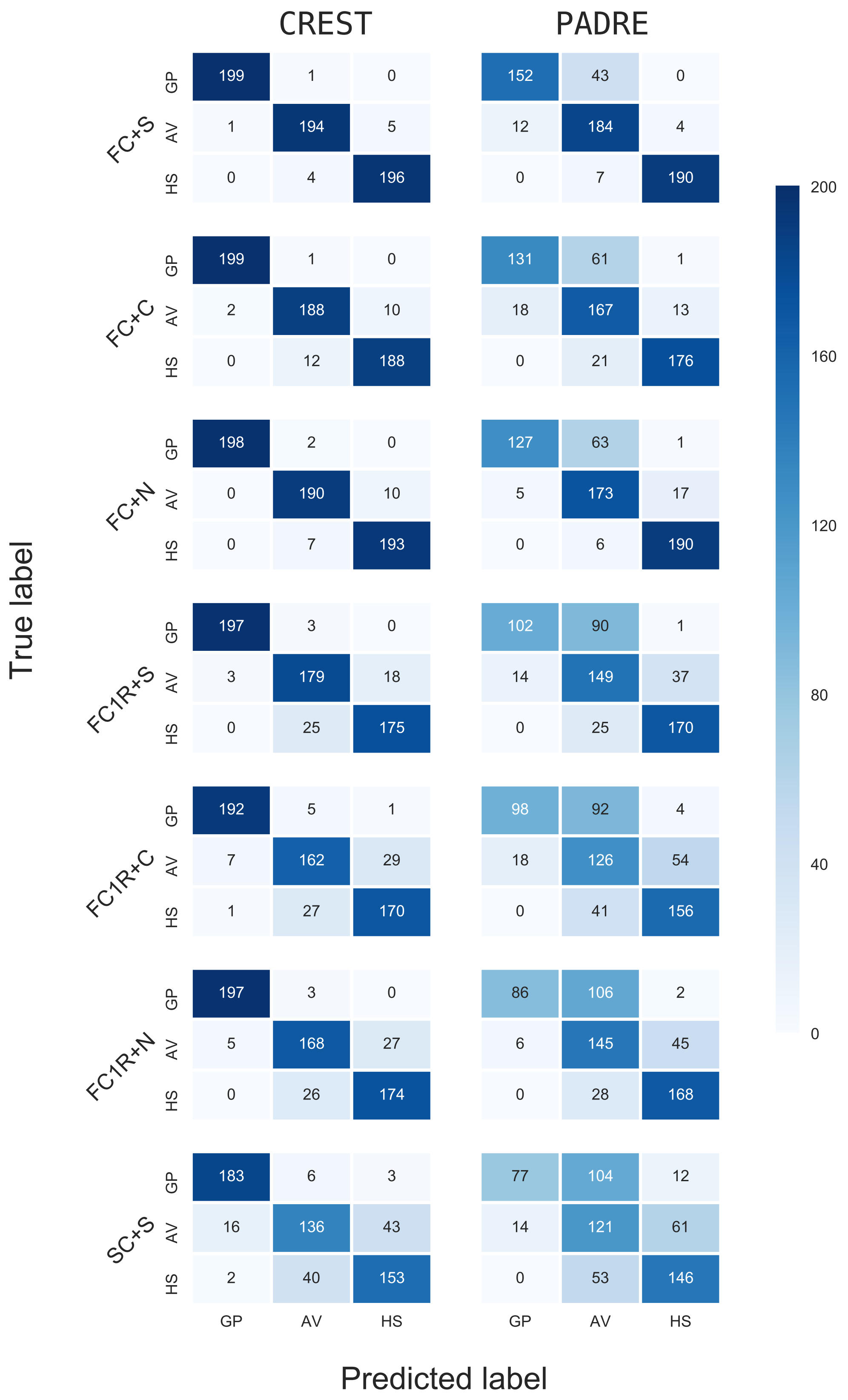


**Figure S4: The confusion matrices from the CREST and PADRE classification results.** Analyses of CREST and PADRE include 200 pairs of GP, AV, and HS over different pedigree structures. Labels on the left indicate the mutual relatives in the pedigree structures. The row of each matrix gives the true relationship type and the column is the predicted relationship type. Since a few pairs failed classification by CREST or PADRE (Results), the sums of each row are not always 200.

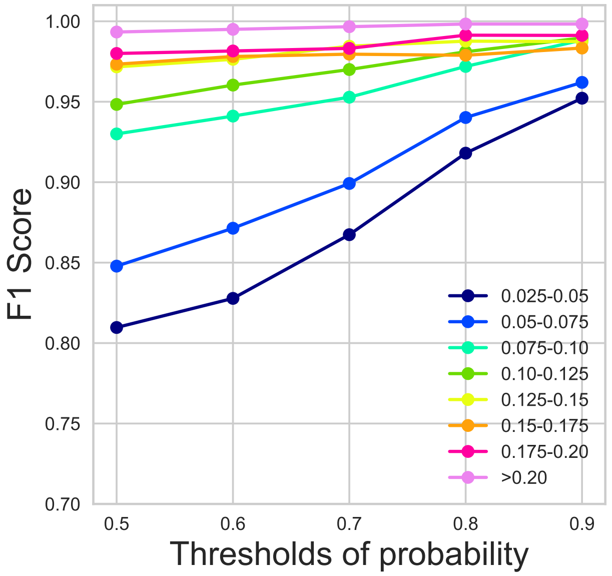


**Figure S5: Thresholding relationship probabilities can improve classification performance.** In every bin of $C$ (depicted as different line colors; legend), we classified relationships using only pairs with a posterior probability above a given threshold and calculated F1 scores from these pairs.

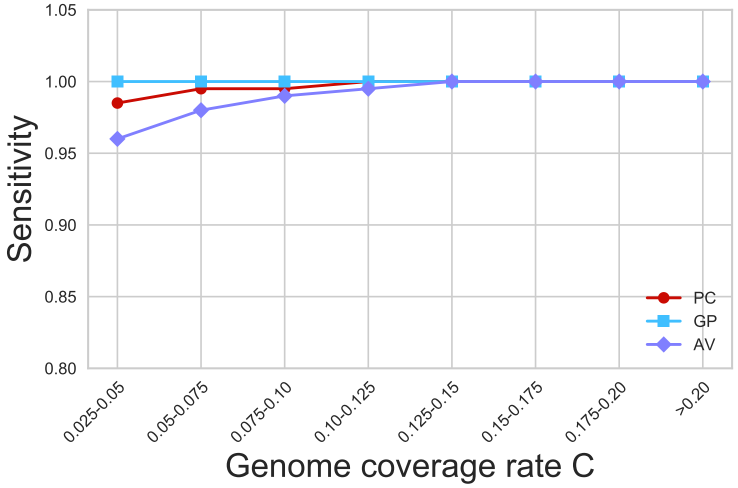


**Figure S6: CREST accurately infers the directionality of PC, GP, and AV pairs.** Plot shows sensitivity across bins of genome coverage rates from other analyses.

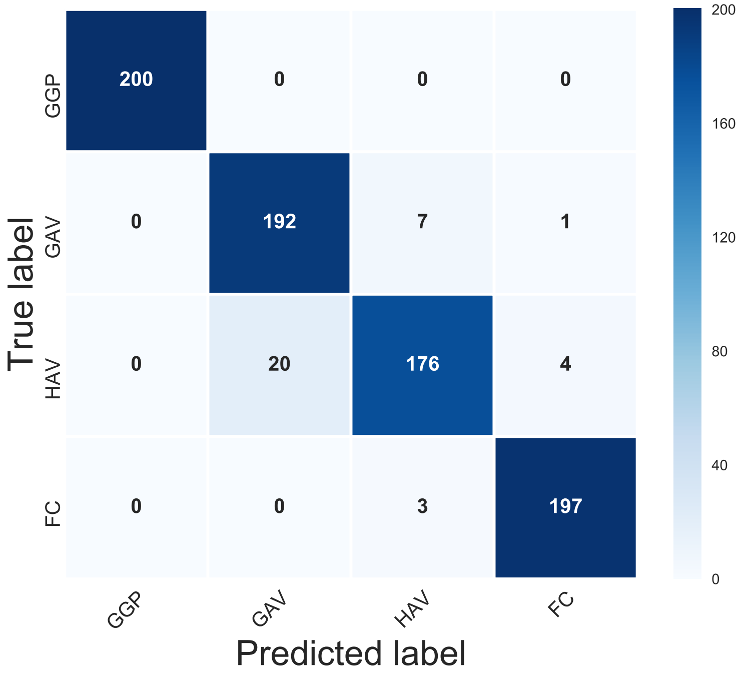


**Figure S7: The confusion matrix for classifying third degree relatives' pedigrees.** The rows correspond to the true relationship type and the column is the predicted type. The analysis includes 200 pairs of each type.

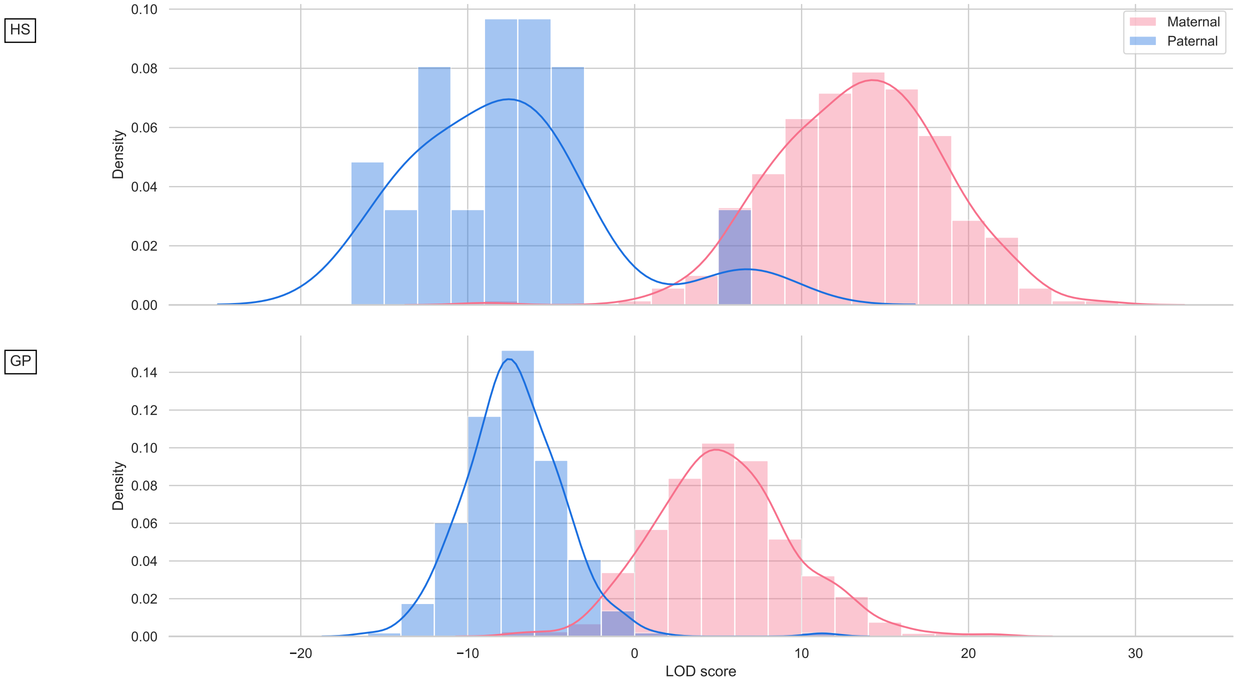


**Figure S8: Initial LOD score histograms from the Generation Scotland data.** Histograms of LOD scores for HS (top) and GP (bottom) pairs include several pairs that are extreme outliers for their reported relationships. We later determined these to be incorrectly labeled in the dataset. Visible anomalies include: a single small maternal HS peak at LOD $\approx$ -8, later determined to be paternal HS; a paternal HS peak including three pairs at LOD $\approx$ 5, later determined to be a maternal HS and two pairs of uncertain relationship; a paternal GP peak at LOD $\approx$ 12, later determined to be an AV pair.
